# OrthoFinder: scalable phylogenetic orthology inference for comparative genomics

**DOI:** 10.1101/2025.07.15.664860

**Authors:** David M Emms, Yi Liu, Laurence Belcher, Jonathan Holmes, Steven Kelly

**Author notes:** Corresponding Author Name: Steven Kelly, Address: Department of Biology, University of Oxford, South Parks Road, Oxford, OX1 3RB, UK.

## Abstract

Here, we present a major advance of the OrthoFinder method. This extends OrthoFinder’s high accuracy comparative genomic framework to provide substantially enhanced scalability and accuracy. Specifically, we show that enhanced phylogenetic delineation of orthogroups provides a 7% relative increase in orthogroup inference accuracy. We further demonstrate that a new gene assignment method substantially reduces overall runtime RAM usage without compromising accuracy. The latest version of OrthoFinder is available at https://github.com/OrthoFinder/OrthoFinder.

## Introduction

Inferring orthology between biological sequences is fundamental to contemporary biological research. It provides the foundation for studying the evolution and diversity of life on Earth and it provides the framework for transfer of biological knowledge between species. Given the central importance of orthology inference to biological research it has been the subject of extensive methodological development for more than 40 years (Fitch, 1970; Gabaldón *et al*., 2009; Altenhoff *et al*., 2016; Linard *et al*., 2021). Diverse approaches to the computational challenge of identifying related genes in different species has resulted in a broad array of methods with varied performance characteristics when applied to diverse datasets (Kristensen *et al*., 2011; Altenhoff *et al*., 2016; Nichio, Marchaukoski and Raittz, 2017).

Although orthology inference has been subject to substantial improvements since the advent of the first automated methods, one of the major challenges facing method development in this field is how to achieve high orthology inference accuracy at scale (Cosentino, Sriswasdi and Iwasaki, 2024; Majidian *et al*., 2025). This challenge has become more pressing in recent years with genome sequencing projects such as The Darwin Tree of Life (The Darwin Tree of Life Project Consortium, 2022) and The Earth BioGenome Project (Lewin *et al*., 2018), which collectively aim to sequence, assemble and annotate the reference genomes for all two million known species of eukaryote. These pioneering efforts, combined with widespread sequencing efforts that are distributed across many biological communities, are creating an unprecedented scalability challenge for automated orthology inference methods (Linard *et al*., 2021; Langschied *et al*., 2024). Thousands of genomes are already available, it is likely that millions more are coming, and there is a real need to be able to accurately and efficiently analyse these resources to maximize their value.

There are several distinct computation bottlenecks for inferring orthology for large numbers of species. For example, the entry point to the majority of inference methods is an all-vs-all sequence similarity search from which orthology relationships can be inferred (Emms and Kelly, 2019). Substantial efforts have been focused at increasing the computational efficiency of individual sequence similarity searches with methods such as DIAMOND (Buchfink, Reuter and Drost, 2021), Usearch (Edgar, 2010), and mmseqs (Hauser, Steinegger and Söding, 2016) greatly advancing the capability to efficiently identify similar sequences in large databases. Even though major improvements have been made, all-vs-all sequence searches have an order of n^2^ time complexity (where n is the number of species under consideration) and consequently are not well suited to conducting orthology inference as species number increases. Thus, alternative orthology inference approaches that retain high accuracy while improving scalability are required to realize the full value of global sequencing efforts.

Previously we developed the OrthoFinder method for phylogenetic orthology inference (Emms and Kelly, 2015, 2019). OrthoFinder first identifies orthogroups, the sets of genes that each descend from a single gene in the last common ancestor of a group of species. OrthoFinder then infers a gene tree for each orthogroup and analyses these gene trees to identify the rooted species tree (Emms and Kelly, 2017, 2018). OrthoFinder also identifies all gene duplication events in the complete set of gene trees and analyses this information in the context of the species tree to provide both gene tree and species tree-level analysis of gene duplication events. Finally, OrthoFinder analyses this cohort of phylogenetic information to identify the complete set of orthologs between all species and provide a suite of comparative genomics statistics. The phylogenetic approach developed in OrthoFinder provided a step change for comparative genomics, enabling the transition from similarity score-based approximations of orthology to tree-based phylogenetic relationships between genes. In recent work, we showed that it was possible to rapidly and accurately place individual sequences into a database of phylogenetic trees and biological sequences produced by an OrthoFinder search (Emms and Kelly, 2022). This approach provided a rapid phylogenetic framework through which individual genes could be added to an existing orthology inference run without incurring a computationally costly all-vs-all re-analysis of existing species. We hypothesized that it should be possible to extend this single-gene approach to support the addition of sets of species and thereby achieve improved scalability without compromising accuracy.

Here we present a major update to OrthoFinder that significantly enhances the scalability of the method. We show that it is possible to rapidly search and assign sequence sets from large numbers of species to a phylogenetically partitioned and structured database of biological sequences in near linear time. We demonstrate that using such an accelerated search functionality does not compromise orthology inference accuracy. Moreover, through advancements in phylogenetic interrogation of orthogroups we show that it is possible to achieve scalable inference with higher accuracy than any competitor method or previous versions of OrthoFinder. Finally, we show that the latest version of OrthoFinder is scalable to thousands of genomes on conventional computing resources. As before, the updated version of OrthoFinder is accurate, customizable, and is performed with a simple command using only protein sequences as input.

## Results

### Improved phylogenetic delineation of orthogroups

In previous versions of OrthoFinder, orthologs were defined phylogenetically but orthogroups were defined solely based on MCL clustering of sequence similarity search results (Emms and Kelly, 2015, 2019). While this method does correct for inter-species divergence it does not leverage the phylogenetic relationship between the species represented in each orthogroup, and thus does not always identify the true extent of an orthogroup. Errors in orthogroup delineation at this stage propagate through all subsequent steps of the analysis, causing collateral errors in orthology inference. To address this, we developed and implemented a novel phylogenetic re-evaluation of orthogroup membership (Figure 1). First, MCL clustering is performed as in OrthoFinder v2, and a gene tree is inferred for each orthogroup. Next, OrthoFinder applies its high accuracy gene tree species tree reconciliation algorithm (Emms and Kelly, 2019) to identify and map all gene duplication events in each gene tree. By definition, an orthogroup should not contain genes arising from ‘ancient’ duplications that predate the root of the species tree, any such duplications indicate that the genes have descended from multiple independent genes present in the last common ancestor of the set of species under consideration. OrthoFinder splits all gene trees at such ancient gene duplication nodes and in doing so creates a new set of revised phylogenetically defined orthogroups which are then recorded and used for all subsequent downstream analyses. OrthoFinder preforms this for all nodes in the species tree creating orthogroups for each node in the species tree. This new step splits erroneously fused orthogroups, and prunes sequences that do not adhere to the phylogenetic definition of the orthogroup (Figure 1).

**Figure 1.**
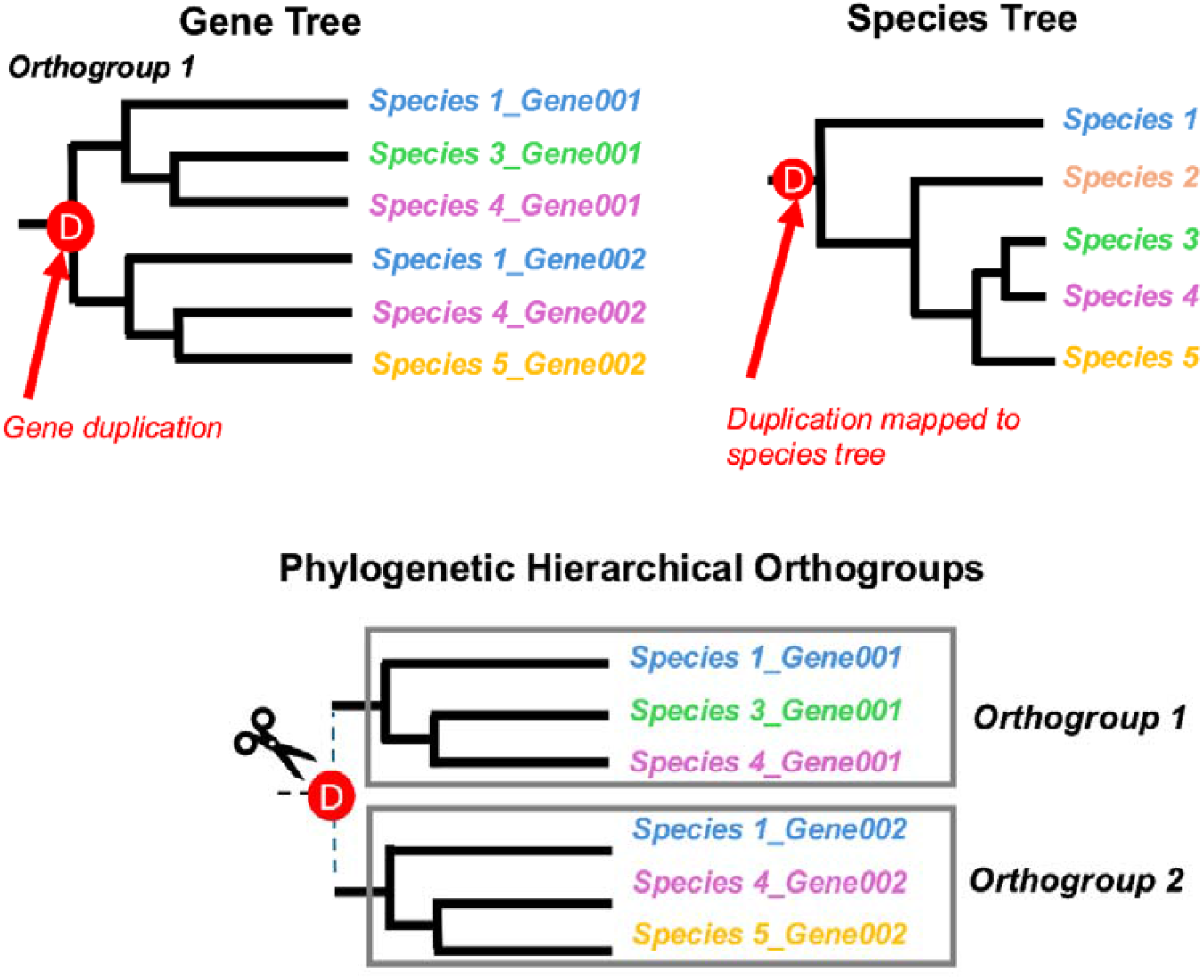
The improved phylogenetic delineation of orthogroups. OrthoFinder reconciles gene trees with the species tree to identify duplication events within putative MCL orthogroups. When a duplication event is identified to have occurred prior to the divergence of the species set under consideration, then orthogroup is split into two (or more) independent orthogroups, ensuring that each resulting orthogroup contains only genes descended from a single gene in the last common ancestor of all species (the definition of an orthogroup).

To test the impact of this novel approach on orthogroup inference accuracy, we compared this new version of OrthoFinder to a set of alternative methods on the OrthoBench dataset of expert-curated reference orthogroups (Trachana *et al*., 2011; Emms and Kelly, 2020). We deployed OrthoFinder in its two most commonly used implementations. One implementation uses multiple sequence alignment and maximum likelihood tree inference to infer orthogroup trees and is provided with a precomputed species tree (OF3_Align_ST), and the other implementation uses the DendroBLAST method to bypass the requirement to construct alignments and alignment-based trees (Kelly and Maini, 2013) (OF3_DB) which is the default implementation in OrthoFinder v2. We also included the new scalable version of OrthoFinder v3 (OF3_Linear) which is discussed in detail in the following section of the manuscript. The list of comparator methods included Broccoli (Derelle, Philippe and Colbourne, 2020), OrthoHMM (Steenwyk *et al*., 2024), SonicParanoid2 (Cosentino, Sriswasdi and Iwasaki, 2024), OrthoMCL (Li, Stoeckert and Roos, 2003), ProteinOrtho (Klemm, Stadler and Lechner, 2023), Hieranoid (Kaduk and Sonnhammer, 2017), and FastOMA (Majidian *et al*., 2025).

For each method, we calculated seven measures of accuracy: (A) the percentage of missing reference orthogroups, (B) the percentage of missing genes, (C) the percentage of incorrect orthogroup fusion events, (D) the percentage of incorrect orthogroup fission events, and (E, F, G) recall, precision, and entropy. Entropy is the measure of reference orthogroup fragmentation, with higher values indicating greater fragmentation. We then calculated an overall rank-based score by evaluating the relative performance of each method across all seven measures. This revealed that OrthoFinder v3 (in either implementation) outperformed all other orthogroup inference methods including OrthoFinder v2 (Figure 2). OrthoFinder v3 had the highest recall, lowest fraction of missing genes, and the lowest entropy. Moreover, OrthoFinder v3 had 5-7% higher accuracy (f-score harmonic mean of recall and precision. Supplementary S1 Table 1) than the previous version of OrthoFinder run with identical settings (OF2_Align_ST and OF2_DB, Figure 2). While OrthoFinder v3 achieved a lower precision than several other methods, OrthoFinder v3 showed a substantially higher recall, and consequently suffered from a much lower rate of missing data. For example, while SonicParanoid2 (SP_def) showed a 3.7% higher precision than OrthoFinder v3 (OF3_Align_ST), the recall of OrthoFinder v3 was 19% higher. Almost all tools were identical in the production of fused reference orthogroups (RefOG), with the exception of FastOMA which erroneously fused 3.5 times more reference orthogroups than other methods (10% compared to 2.85%). The spread of reference orthogroup fissions events across the tools varied, with low precision tools such as Broccoli and OrthoHMM making relatively few RefOG fissions. Analysis of the entropy scores, which are similar to a weighted combination of fission and fusion events, revealed that OrthoFinder tends to produce orthogroups whose sizes and membership most accurately recapture the reference dataset (Figure 2). Therefore, the introduction of the phylogenetic delimitation of orthogroups improves overall orthogroup inference accuracy, and OrthoFinder v3 is the most accurate method currently available.

**Figure 2.**
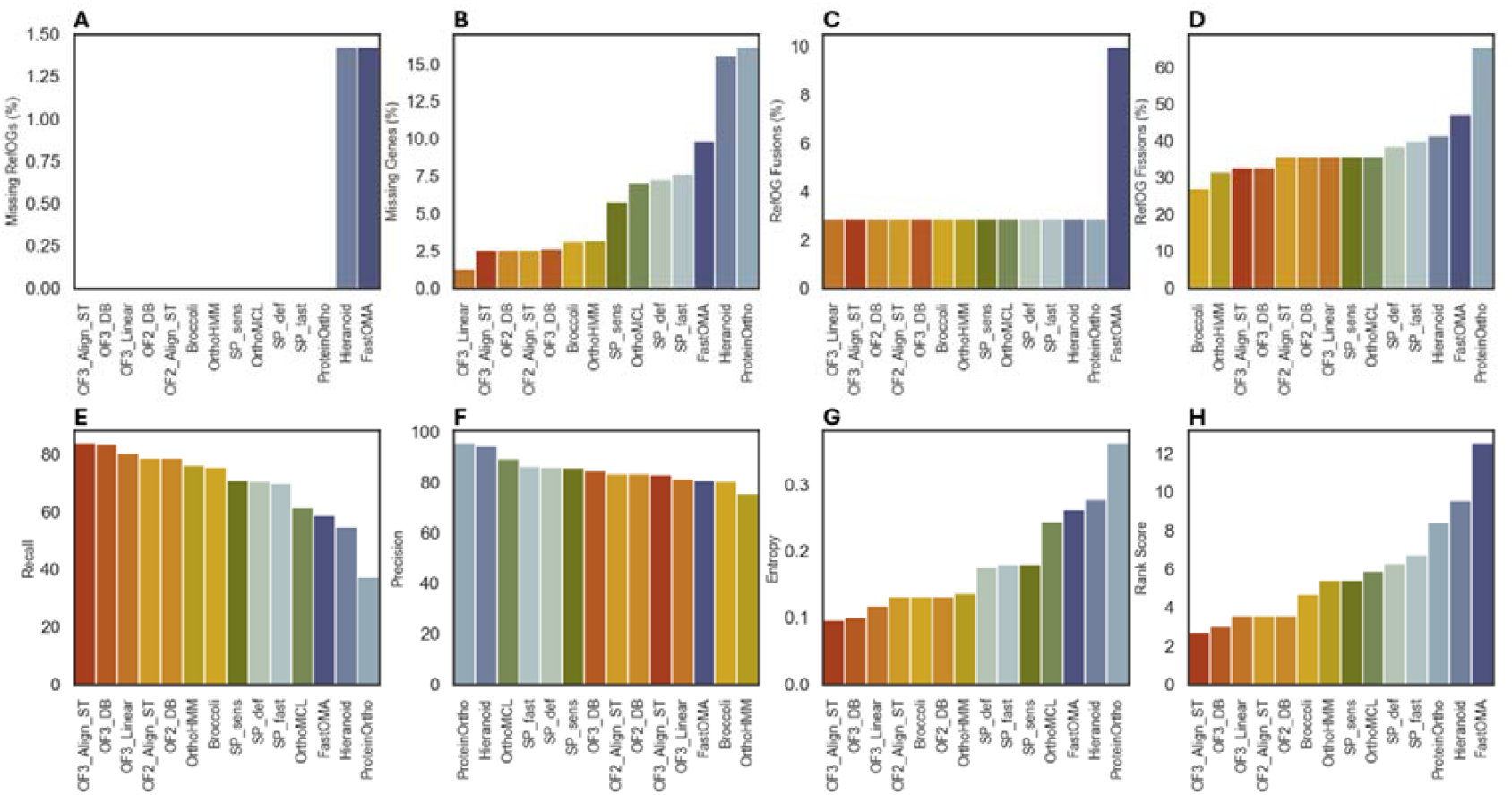
Comparative analysis of different orthogroup inference tools on the OrthoBench dataset. Panels A-G each show one of seven performance metrics used to evaluate orthology prediction accuracy. Lower values indicate better performance for error-based metrics (A-D,G), higher values are better for recall and precision (E-F). Panel H shows the rank score, combining all seven metrics.

### Achieving enhanced scalability without compromising accuracy

In addition to improvements in the phylogenetic definition of orthogroups, we also developed a novel implementation of OrthoFinder to improve the scalability of the method. This is achieved through adaptation and further development of the SHOOT profile algorithm (Emms and Kelly, 2022) which enables rapid addition of new species to existing OrthoFinder runs. This novel scalable version of OrthoFinder is implemented as a two-step process (Figure 3). Prior to starting, the input species set are partitioned into two non-overlapping subsets – a “core” subset and an “assign” subset. It is recommended that the “core” subset contain fewer than 100 species for analysis on conventional computing resources. The “assign” subset can be substantially larger as described below. The first step of this new implementation requires that a conventional OrthoFinder run is performed on the core subset. This creates a phylogenetically partitioned and structured reference database of biological sequences (Emms and Kelly, 2022) that is then used as input for the second stage of the algorithm. The species from the assign subset are then rapidly assigned to the correct orthogroup in the reference database, irrespective of the number of sequences contained within each core orthogroup. Following completion of the assignment step, OrthoFinder’s rapid phylogeny-based orthogroup and ortholog inference steps are performed to produce an expanded rooted species tree, gene trees, phylogenetically determined orthogroups, orthologs, gene duplication events, and comparative genomic statistics (Figure 3).

**Figure 3.**
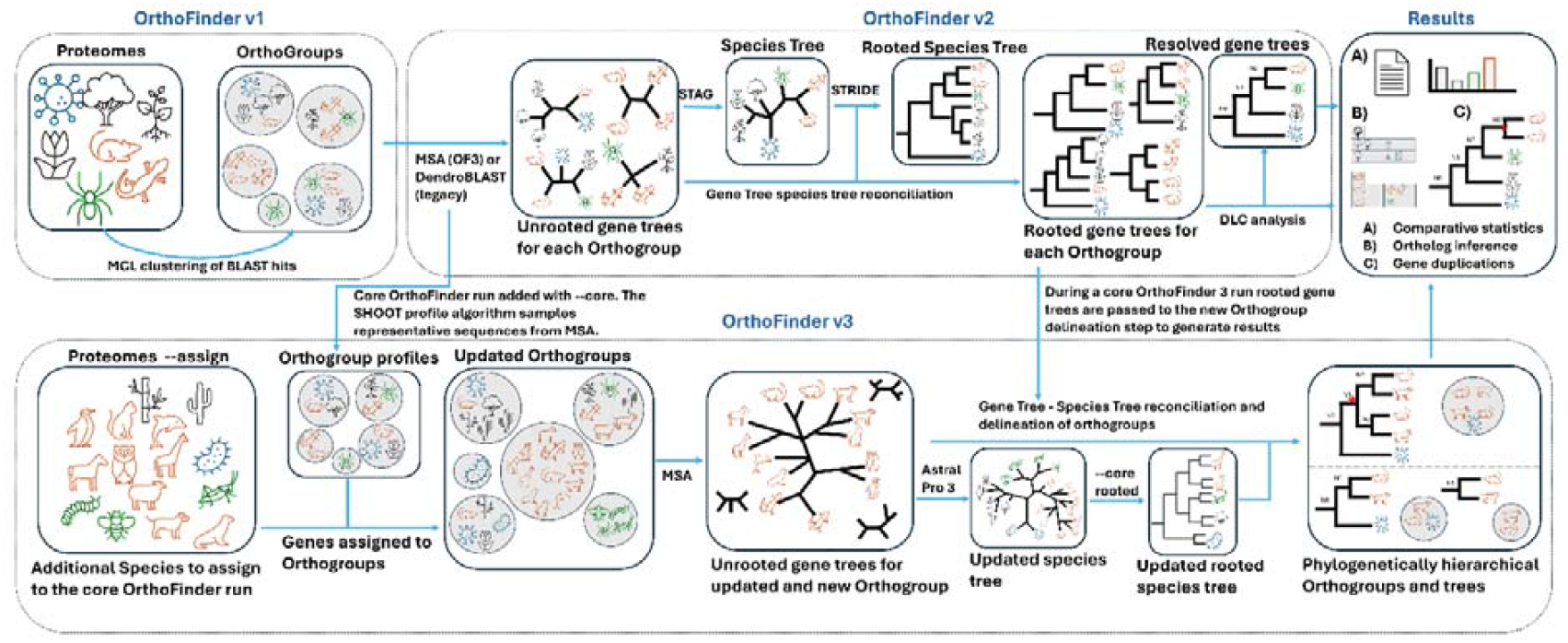
The OrthoFinder v3 workflow. The first step of OrthoFinder 3 is to perform a conventional OrthoFinder run on a set of core proteomes. The second step then uses an adapted version of the SHOOT profiling algorithm to assign genes from additional species to these core orthogroups and identify new orthogroups not present in the initial input gene set. Gene trees are then inferred and used to infer a revised species tree. All results files are then revised and updated.

To test the impact on scalability of this new workflow, OrthoFinder v3 was compared against the set of commonly used orthology inference tools. The time each method took to complete analyses of datasets of different sizes was measured, unless a timeout of 7 days was reached before completion. Peak memory usage was also calculated for each analysis. Each tool was run on a Linux server using AMD processers, all tools were allocated 32 threads (16 cores) and up to 200gb of RAM for each run. In order to test performance on a representative set of species, the Ensembl (Harrison *et al*., 2024) rapid release genomes (Accessed 29^th^ of August 2024) were downloaded. Test datasets ranging in size from 2 - 1024 species we compiled by sampling species from Ensembl dataset using PDA (phylogenetic diversity analyzer) (Chernomor *et al*., 2015), see ‘Methods’ for more details. All methods were run on the same datasets in their default implementations unless indicated otherwise in the methods or figure legends. OrthoFinder v3 was run using the standard OrthoFinder 2 workflow up to 64 species. Beyond 64 species OrthoFinder v3 was run in the new scalable implementation where the first 64 species were taken to be the core set and all other species were assigned. The final run time for OrthoFinder v3 was calculated using the combined run time for the core 64 species and the assigned dataset.

OrthoFinder v3 linear addition outperformed all other tested methods, being the only method able to perform orthology inference on 1024 species within the seven-day cutoff (Figure 4A). OrthoFinder v3 completed this run in 128 hours. SonicParanoid2 (fast mode) and FastOMA were the only two methods able to run 512 proteomes within the cutoff time. OrthoFinder v3 followed a near linear run time trend for 128 – 1024 species. This new implementation of OrthoFinder was faster than each of the other competitor methods tested and significantly (x8) faster than OrthoFinder v2 on datasets larger than 64 species, an important improvement in scalability from the previous version.

**Figure 4.**
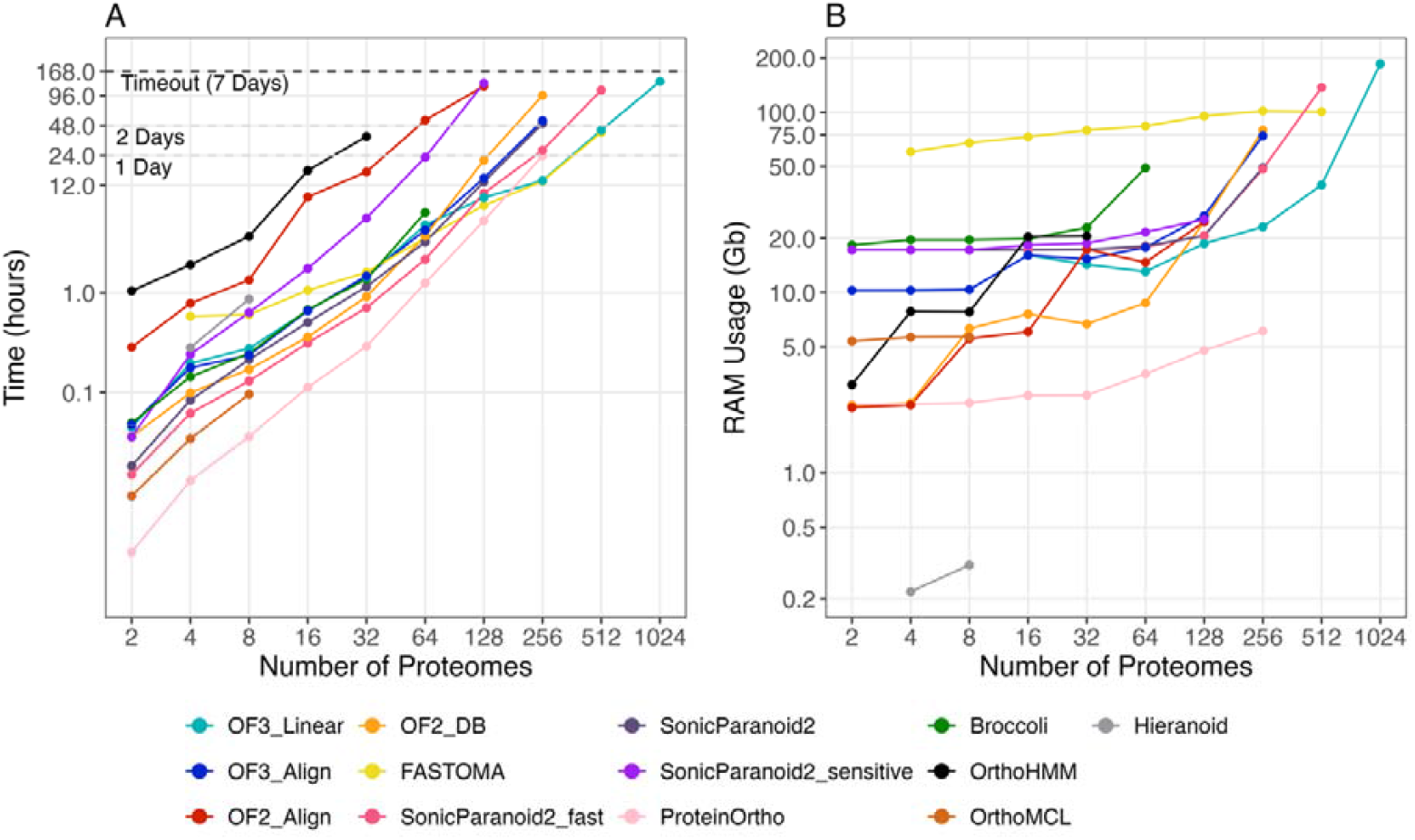
A comparison of runtime and peak memory usage for orthogroup inference tools. (A) Time (log scale) and (B) peak RAM usage required to run different orthology prediction tools across increasing numbers of input proteomes (from 2 to 1024).

OrthoFinder v3 also performed well on memory usage (Figure 4B). OrthoFinder v3 outperformed OrthoFinder v2 above 128 species, with increasing memory saving as species number increases. For example, v3 linear shows a 3.4-fold decrease in RAM consumption for 256 species compared to v2 DendroBlast. On the largest datasets that other methods could complete, OrthoFinder v3, used approximately 4-fold lower peak memory consumption (Figure 4B).

To demonstrate that the alternative implementation did not compromise orthogroup inference accuracy, the linear species addition method (OF3_Linear) was also tested on the OrthoBench data (Figure 2). OrthoFinder v3 Linear only slightly underperformed compared to the non-scalable version, with slight decreases in some measures observed. However, this scalable version was more accurate than any implementation of OrthoFinder v2 and any other competitor methods (Figure 2), supporting the use of this accelerated implementation of OrthoFinder in comparative genomic analyses.

### OrthoFinder 3 performs accurate ortholog inference on the quest for orthologs benchmarking service

We also tested the accuracy of the orthologs predicted by OrthoFinder v3 in multiple implementations using the Quest for Orthologs (QfO) benchmarking service (Gabaldón *et al*., 2009; Altenhoff *et al*., 2016; Linard *et al*., 2021). QfO is the most widely used orthology benchmarking service, which evaluates a tool’s ability to accurately predict orthologs across a diverse set of taxa, including Archaea, Bacteria, and eukaryotes. QfO gives many different measures of accuracy, including comparisons to expert-curated reference orthologs and the accuracy of species trees inferred from a set of orthologs. We used the most recent published set of reference proteomes, which consists of 78 species (23 bacteria, 48 eukaryotes, and 7 archaea) (Nevers *et al*., 2022). We compared OrthoFinder v3 against the results obtained from 14 orthology inference tools.

Analysis of the resulting scores revealed that the new OrthoFinder v3 workflow for high scalability does not compromise accuracy (Figure 5). OrthoFinder v3 is on the Pareto frontier for species tree discordance tests in both eukaryotes (Figure 5A), and bacteria (Figure 5B), demonstrating its ability to perform accurate orthology inference across different domains of life. The only other tool with scalability comparable to OrthoFinder v3 is FastOMA, but this method has substantially lower ortholog inference accuracy (Figure 5). For example, while the Robinson-Foulds distance (measuring disagreement between true and inferred species trees) for eukaryotes is marginally worse in OrthoFinder v3 compared to FastOMA (0.06 vs. 0.05), OrthoFinder v3 has an 80% higher recall (15721 vs. 8686). Similarly, OrthoFinder v3 is marginally better than FastOMA for Robinson-Foulds distance on the bacteria species tree discordance test (0.590 vs. 0.587), but again has a 23% higher recall. OrthoFinder v3 is on the Pareto frontier for the enzyme classification test (Figure 5C), which measures how well predicted orthologs match to Enzyme Commission numbers (which are given depending on the chemical reaction they catalyze). OrthoFinder v3 is only outperformed in precision by tools that rely on pre-computed databases (e.g. OrthoInspector and PANTHER), and is better than FastOMA in both precision (0.933 vs 0.928) and recall (183368 vs 157049).

**Figure 5.**
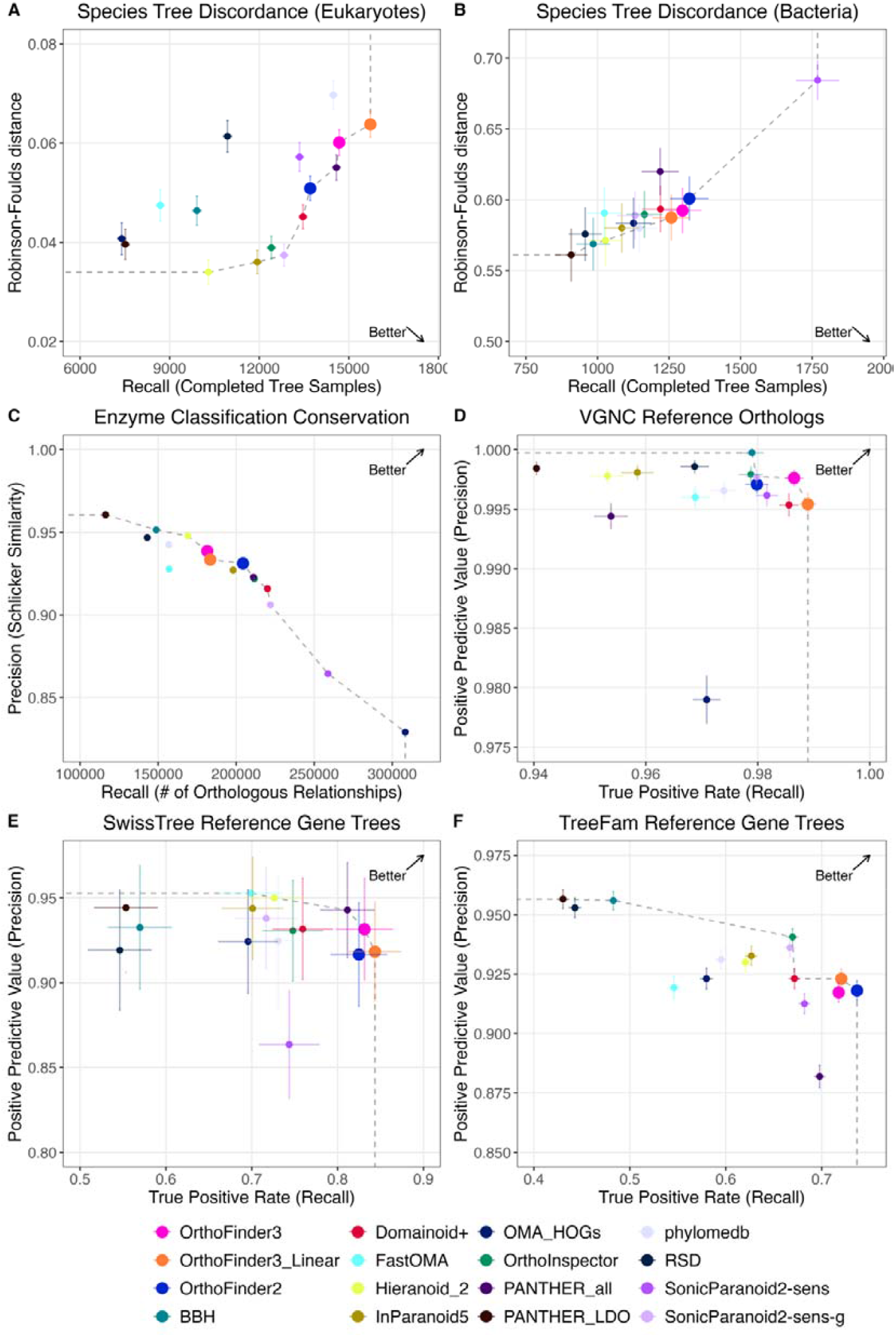
A comparative analysis of ortholog inference accuracy on the Quest for Orthologs dataset. Each panel shows a different benchmark from the QfO benchmarking service (2022 release). OrthoFinder is compared against other ortholog inference methods which have made their results publicly available. (A–B) Species tree discordance (Robinson-Foulds distance) vs. recall across completed gene trees for eukaryotes (A) and bacteria (B). (C) Conservation of enzyme classification across orthologs (Schlicker similarity) vs. number of orthologous relationships. (D) VGNC reference orthologs: precision vs. recall. (E–F) SwissTree (E) and TreeFam (F) reference gene trees: precision vs. recall. Better-performing methods are toward the top-right for precision measures (C-F), and bottom right for discordance measures (A-B).

OrthoFinder v3 is on the Pareto frontier for all three of the human-curated reference sets (Figure 5 D-F), scoring particularly well on recall. OrthoFinder v3 has the highest recall of all methods on both the VGNC and SwissTree human-curated reference sets and is only marginally beaten for recall on the TreeFam dataset by OrthoFinder v2 (0.72 vs. 0.74). In summary, similar to the high accuracy of its orthogroup inference, OrthoFinder v3 also exhibits high accuracy ortholog inference across multiple benchmark tests.

## Discussion

Planet Earth is home to more than 6,000 species of mammal (Burgin *et al*., 2018), 300,000 species of plant (Christenhusz and Byng, 2016), 5,000,000 species of insect (Gaston, 1991), and an unknown number of species of unicellular eukaryotes, bacteria and archaea (Mora *et al*., 2011; Hugenholtz *et al*., 2021). Inferring the phylogenetic relationships between the biological sequences of these organisms provides a foundation upon which we can study evolution and molecular diversity, and enables us to understand and transfer biological information between organisms. It is likely that a substantial fraction of all known species on Earth will have a representative genome in the coming decades, but methods do not yet exist to enable this scale of data to be analysed. Here, we present a major advance to OrthoFinder that substantially improves the scalability and accuracy of the method. We show it is possible to rapidly assign gene sets from large numbers of species to a phylogenetically partitioned and structured database of biological sequences in near linear time. We also show that using phylogenetic delimitation improves the inference of orthogroups, enhancing the accuracy of OrthoFinder and extending its performance advantage over comparator methods. OrthoFinder remains an easy-to-use, fast, accurate, and fully phylogenetic orthology inference software tool. OrthoFinder also continues to be one of the few tools to provide a wide range of output information including gene duplications, gene trees, sequence alignments and single copy ortholog sequences allowing downstream analysis.

The updated OrthoFinder method is now capable of analysing thousands of species using a novel two-step process. Although this process requires an additional step for the user, the only input required is still the set amino acid sequences corresponding to the protein-coding genes for the species of interest. The default parameters for OrthoFinder have been optimized for speed, accuracy, and scalability and enable the combined analysis of thousands of species on commonly available computing resources. OrthoFinder also retains its customizability for expert users, and intermediate steps in the algorithm (such as alignment or tree inference) can be substituted with alternative methods should the user wish. Although there is still some distance to travel, this upgrade to OrthoFinder provides an important step towards the goal of being able to provide high accuracy phylogenetic orthology inference for all species on Earth.

## Materials and Methods

### The OrthoFinder v3 workflow enhanced scalability workflow

The scalable linear species addition framework of OrthoFinder v3 requires the results of a conventional OrthoFinder analysis of an ‘core’ set of species. For each orthogroup, OrthoFinder v3 employs the SHOOT profile algorithm (Emms and Kelly, 2022) to select the most representative sequences using k-means clustering applied to an embedding of the sequences from the length L multiple sequence alignment (MSA) of the orthogroup in a 2*L-dimensional space. DIAMOND (Buchfink, Reuter and Drost, 2021) is used to assign the new genes to the core orthogroups using these profiles. Genes not assigned to any core orthogroups are set aside to be analysed separately at a later stage. Once all genes have been assigned, gene trees are inferred. To support the analysis of larger datasets, the MAFFT multiple sequence alignment method has been replaced with the more scalable FAMSA method (Deorowicz, Debudaj-Grabysz and Gudyś, 2016). Users can revert back to previous default alignment method “-A mafft”. Maximum likelihood tree inference of the resulting alignments is then performed using FastTree (Price, Dehal and Arkin, 2010) employing the “-fastest” runtime option. The use of other alignment or tree inference programs is supported through use of the config.json file, although users are advised to test if they have sufficient computational resources to use more computationally expensive programs. Once all orthogroup trees are inferred, a revised species tree is then computed using ASTRAL-Pro (Zhang and Mirarab, 2022).

As mentioned above, some genes fail to be assigned to any core orthogroups during the initial assignment step. This occurs because the new species (or sets of species) being added will sometimes contain new genes that have arisen *de novo*, and are not found in the core species set. To resolve this, OrthoFinder performs a root to tip tree traversal to identify clades of species (minimum size = 2) that had no representation in the now sparsely sampled initial core species set. Once identified a conventional OrthoFinder analysis is performed on the clade, and the new orthogroups are added to the revised orthogroup set.

The original OrthoFinder algorithm applied a per-species-pair gene length normalisation that accounted for both evolutionary-distance and gene-length dependence in pairwise bit-scores between sequences (Emms and Kelly, 2019). However, this normalisation step requires a large number of pairwise hits between the genes in each species in order to fit the required correction functions. To enable the smaller set analysis above, the updated version of OrthoFinder employs an alternative normalisation approach. The approach separates the sequence length bias and species divergences to obtain many independent estimates of the extent of each orthogroup Specifically, bit-scores between two sequences, B_qh_, are length normalised using geometric mean of the lengths of the query and hit sequences, L_q_ and L_h_: B_qh_=B_qh_/(L_q_L_h_)^1/2^. As before, normalised reciprocal best hits (RBH) are treated as putative orthologs and are used to estimate the sequence divergence between pairs of species and on a per-orthogroup basis. In the modified version we calculate a set of ratios R_XY_* for species X and Y. These estimate the expected ratio of the normalised bit score between the RBH for a pair of genes x and y from species X and Y from some orthogroup and the normalised bit score from the gene x to the most distant gene in that orthogroup. Using these, every RBH observed can be used to estimate a cut-off for the most distant gene in that orthogroup. In this way, if a gene has an RBH in multiple species then it provides multiple independent estimates of the cut-off for the most distant sequence in the orthogroup. Likewise, if multiple genes from a (currently undetermined) orthogroup obtain one or more RBH, then these will each provide multiple independent estimates of the extent of the orthogroup. These estimates are used to construct a graph such that a pair of sequences is connected within the graph if the NBS between those sequences fall within that gene-specific and species-specific cut-off. As such, this graph attempts to connect as densely as possible with edges all sequences within the same orthogroup while having as few edges as possible between sequences in different orthogroups. As before, the MCL algorithm (Van Dongen, 2008) is applied to identify the sets of genes in the graph best satisfying the putative shared orthogroup memberships encoded in the edges of this graph. As with the core orthogroups, the resulting gene trees of the putative orthogroups are analysed to phylogenetically resolve the true extent of each orthogroup (Figure 1). The non-core orthogroups are subjected to phylogenetic analysis to infer rooted gene trees and subsequently these rooted gene trees are analysed to determine the full comparative analysis including orthologs, hierarchical orthogroups, and gene duplication events.

### Assessing orthogroup inference accuracy

The full details on the orthogroup benchmarking, including the calculation of the scores, and a complete method by method breakdown of the results are provided in Supplemental File 1 and the complete set of output files generated by each method is provided in Supplemental File 2.

### Assessing scalability performance

The Ensembl (Harrison *et al*., 2024) rapid release dataset was accessed on the 29^th^ August 2024. Amino acid sequences for all protein coding genes in each genome were downloaded. For species which had more than one version available, the most recent proteome file was used. The OrthoFinder v2 script ‘primary_transcript.py’ was used to extract only the longest variant for each gene giving a dataset with a total of 1789 species. Following collection of the Ensembl proteomes an approximate species tree was derived in order to extract a set of representative proteomes for performance testing across each orthology inference tool. The species tree was generated using orthologs of BUSCO genes (Manni *et al*., 2021). In brief, the hidden Markov model provided for each BUSCO gene was used to identify high scoring amino acid sequences in each species. The resulting hits from each species to each BUSCO gene were aligned using MAFFT, and the resulting multiple sequence alignments concatenated. An unrooted phylogenetic tree was then inferred from this concatenated alignment using FastTree. This tree was then used as input to Phylogenetic Diversity Analyzer (Chernomor *et al*., 2015) to select sets of species in the range of 2-1024 species such that each subsequent set contained all the species of the previous set, and that each subset maximised the sampled branch length of the inferred tree. The complete result dataset for runtime and scalability of all methods is provided in Supplemental File 3.

### Assessing ortholog inference accuracy

OrthoFinder v3 was run on the 2022 edition of the Quest for Orthologs reference proteomes. We ran both OrthoFinder v3 Align (with FAMSA), and OrthoFinder v3 Linear (−-core –assign). OrthoFinder v2 along with 13 other orthology inference tools have already been run on this dataset, and is available as part of the publicly available results alongside many methods. The Quest for Orthologs datasets consists of 78 proteomes (48 eukaryotes, 23 prokaryotes, and 7 archaea). We ran OrthoFinder on these input datasets and extracted pairs of orthologous genes identified by the method. The file containing orthologous pairs was uploaded to the Quest for Orthologs web server for benchmarking. The Quest for Orthologs papers describe the methodology for benchmarking in full detail (Altenhoff *et al*., 2016; Nevers *et al*., 2022).

## Acknowledgements

The authors would like to thank the OrthoFinder user community for their feedback on the method and their continued support.

## Author information

DE developed the method. DE, YL, LB, JH implemented software. DE, YL, LB, JH, SK contributed to the analysis. DE, LB, JH, SK wrote and edited the manuscript.

## Ethics declaration

Sk is co-founder of Wild Bioscience Ltd and an employee of Ellison Institute of Technology, Oxford Limited.

