## Supplemental File 1 for "OrthoFinder: scalable phylogenetic orthology inference for comparative genomics"

Supplement S1: Orthogroup Benchmarking

The aim of orthogroup benchmarking is to assess the accuracy of the set of orthogroups predicted by an orthology inference tool. We do this by comparing a set of predicted orthogroups to the ‘OrthoBench’ reference set of expert-curated orthogroups (Emms et al. 2020).

The OrthoBench data benchmarking problem can be mathematically summarised as follows.

Let  $\mathcal{G=}\left\{ g_{1},g_{2}, \ldots,g_{N} \right\}, N\mathbb{\in N}$ be a countable finite set of genes of interest. Given two clustering of $\text{R = }\left\{ R_{1},R_{2}, \ldots,R_{S} \right\}\text{ }$ with $S\mathbb{\in N} , \bigcup_{i=1}^{S} R_{i}\mathcal{\subseteq G}$ and $\text{P = }\left\{ P_{1},P_{2}, \ldots,P_{L} \right\}$ with $L\mathbb{\in N,}\bigcup_{j=1}^{L} P_{i}\subseteq\mathcal{G}$), they represent the reference orthogroups (RefOGs) and predicted orthogroups (PredOGs), respectively.

The partitions in RefOGs and PredOGs are pairwise disjoint, i.e., $\bigcap_{i=1}^{S} R_{i}=\bigcap_{j=1}^{L} P_{i}=\emptyset$. The goal of benchmarking is to quantify the difference between the two clusterings $\text{R}$ and $\text{P}$. Here, we proposed seven measures to achieve this goal.

1. Missing RefOGs (%)

Summary: Missing RefOGs (%) measures the percentage of reference orthogroups that have no genes recovered in any predicted orthogroup.

In certain methods, it is possible for no genes to be identified in a partition derived from reference orthogroups. This means that $\exists M\mathbb{\in N,}M<S$, $\sum_{i=0}^{M} \sum_{j=0}^{L} \left| R_{i}\bigcap P_{j} \right|=0$. In other words, $\exists\boldsymbol{MR\subseteq R}$, such that

$$\mathbf{MR}=\left\{ R_{i}\in\text{R } | \forall P_{j}\in\text{P }{, R}_{i}\cap P_{j}=\emptyset\right\}$$

With this definition, the missing RefOGs (%) is given by

$$100\times\frac{\left| \mathbf{MR} \right|}{S}$$

where $\left| \mathbf{MR} \right|\boldsymbol{=}M$ counts the number of missing RefOGs.

2. Missing Genes (%)

Summary: Missing Genes (%) measures the percentage of all genes in reference orthogroups that are not found in any predicted orthogroup.

As for the missing genes, we can first define a missing genes set for each RefOG, namely

$$\mathbf{MG}\left( R_{i} \right)= \left\{ g\in R_{i} | \forall P_{j}\in\mathbf{P,}g\notin P_{j} \right\}$$

Then the missing genes measure can be computed as

$$100\times\frac{\sum_{i=0}^{S} \left| \mathbf{MG}\left( R_{i} \right) \right|}{S}$$

where $\left| \mathbf{MG}\left( R_{i} \right) \right|$ is the size of the missing genes set of a RefOG $R_{i}$.

3. RefOG Fusions (%)

Summary: RefOG Fusions (%) measures the percentage of reference orthogroups that have instances where they are merged into the same predicted orthogroup as another reference orthogroup. This indicates over-clustering of orthogroups.

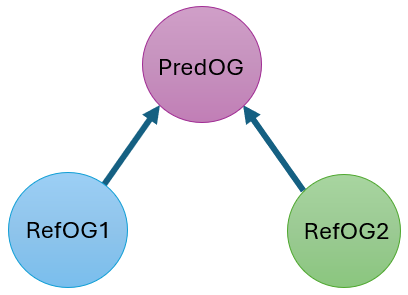

This measure quantifies the number of RefOGs that are associated with a shared PredOG. It captures situations where multiple RefOGs overlap with the same PredOG, we define

$$\mathbf{R}^{'}\left( P_{j} \right)=\left\{ R_{i}\in\mathbf{R} | R_{i}\bigcap P_{j}\neq\emptyset\right\}$$

which represents a set of RefOGs that have non-empty intersection with $P_{j}$. Then $\forall P_{j}\in\text{P}$, the union of the $\mathbf{R}^{'}\left( P_{j} \right)$ when $\left| \mathbf{R}^{'}\left( P_{j} \right) \right|>1$ gives the all the RefOGs that have shared PredOGs, can be rewritten as

$$\mathbf{R}^{'}= \bigcup_{\forall P_{j}\in\text{P: }\left| \mathbf{R}^{'}\left( P_{j} \right) \right|>1} \mathbf{R}^{'}\left( P_{j} \right)$$

Then the fusion score can be obtained by dividing the cardinality of $\mathbf{R}^{'}$ (i.e., the number of fused RefOGs) over all the total number of RefOGs, namely

$$100\times\frac{\left| \mathbf{R}^{'} \right|}{S}$$

4. RefOG Fissions (%)

Summary: RefOG Fissions (%) measures the percentage of reference orthogroups that have instances where they are split across multiple predicted orthogroups. This indicates over-fragmentation of orthogroups.

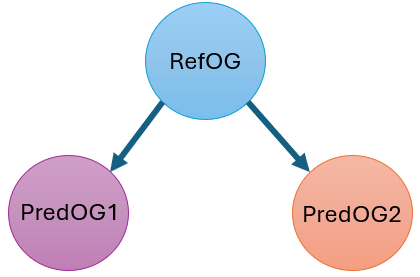

As the opposite of Fusion, Fission captures percentage of RefOGs are split into multiple PredOGs, which can be defined as follows:

$$100\times\frac{\sum_{i=0}^{S} \boldsymbol{1}_{\left| \mathbf{P}^{'}\left( R_{i} \right) \right|>1}}{S}$$

where

$$\mathbf{P}^{'}\left( R_{i} \right)= \left\{ P_{j}\in\mathbf{P} | R_{i}\bigcap P_{j}\neq\emptyset\right\}$$

Represents the set of PredOGs that overlap with a given RefOG $R_{i}$, and the indicator function evaluates to 1 only when $\left| \mathbf{P}^{'}\left( R_{i} \right) \right|>1$, i.e., $R_{i}$ overlaps with more than one PredOG, indicating a fission.

5. Recall

Summary: Recall measures the proportion of genes in each reference orthogroup that are recovered in predicted orthogroups, computed as a weighted average across overlaps and then averaged over all reference orthogroups with size-based weighting.

Recall and Missing Genes (%) are complementary metrics that both evaluate the recovery of genes from reference orthogroups. Recall emphasizes the proportion of successfully identified genes, while Missing Genes (%) highlights gene loss.

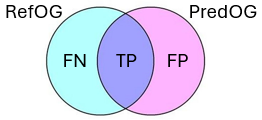

As a well-known measure, recall measures how well a method identifies all relevant instances (true positives) from the total number of actual relevant instances, which is defined as follows:

$$RC= \frac{TP}{TP+FN}$$

where TP stands for true positives, FN stands for false negatives.

Following this definition, the recall between the $i$th RefOG and the $j$th PrefOG is given by

$${rc}_{ij}=\frac{\left| R_{i}\bigcap P_{j} \right|}{\left| R_{i} \right|}$$

where $\left| R_{i}\bigcap P_{j} \right|$ is the size of the intersection between RefOG $R_{i}$ and PredOG $P_{j}$. $\left| R_{i} \right|$ is the size of RefOG $R_{i}$.

Since a RefOG can be split into multiple PredOGs, in this study, we favor the weighted average version of the recall score to fully capture the relationship between the RefOGs and the PredOGs. For each RefOG, we can define the weighted average recall as follows:

$${RC}_{i}=\frac{\sum_{j=0}^{L} \left| R_{i}\bigcap P_{j} \right|*{rc}_{ij}}{\sum_{j=0}^{C} \left| R_{i}\bigcap P_{j} \right|}$$

Under this context, $\left| R_{i}\bigcap P_{j} \right|$ is chosen as the weight.

To obtain the total average scores for all RefOGs, we can apply the same idea across all RefOGs to get the total weighted average recall score, namely,

$$\mathbf{RC}= \frac{\sum_{i=0}^{S} \left| R_{i} \right|*{RC}_{i}}{\sum_{i=0}^{S} \left| R_{i} \right|}$$

Here, the size of each RefOG $\left| R_{i} \right|$ is used as the weight.

6. Precision

Summary: Precision measures the proportion of genes in each predicted orthogroup that correctly match genes from the corresponding reference orthogroup, computed through overlap-based weighting and averaged across all reference orthogroups.

For precision, it’s general definition is given by the following expression,

$$PS= \frac{TP}{TP+FP}$$

where $FP$ stands for false positive. Then, the precision between the $i$th RefOG and the $j$th PrefOG is given by

$${ps}_{ij}=\frac{\left| R_{i}\bigcap P_{j} \right|}{\left| P_{j} \right|}$$

Similarly we can define the weighted average precision for each RefOG and the total weighted average precision for all RefOGs as

$${PS}_{i}=\frac{\sum_{j=0}^{L} \left| R_{i}\bigcap P_{j} \right|*{ps}_{ij}}{\sum_{j=0}^{C} \left| R_{i}\bigcap P_{j} \right|}$$

and

$$\mathbf{PS=}\frac{\sum_{i=0}^{S} \left| R_{i} \right|*{PS}_{i}}{\sum_{i=0}^{S} \left| R_{i} \right|}$$

respectively.

7. Entropy

Summary: Entropy measures the fragmentation of each reference orthogroup by computing the Shannon entropy of how its genes are distributed across predicted orthogroups (including an extra category for missing genes), with the final score being the size-weighted average of normalized entropy values across all reference orthogroups — higher values indicate greater dispersion of genes across predictions.

Both RefOG Fission (%) and Entropy assess how genes from a reference orthogroup are distributed across predicted orthogroups; Fission provides a coarse, binary indicator of splitting events, whereas Entropy offers a fine-grained, Shannon entropy–based measure of fragmentation severity.

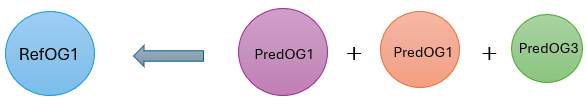

Since each RefOG can be split into multiple PredOGs, we introduce entropy as a measure to capture how spread out such effect is.

For each cluster in $\mathbf{R}$, each gene can be assigned a different label if it is classified into different PredOGs. Therefore, we define the set

$$\mathbf{C}\left( R_{i} \right)= \left\{ P_{j}\in\mathbf{P} | R_{i}\bigcap P_{j}\neq\emptyset\right\}\bigcup\left\{ P_{m} | g\in R_{i}, g\notin\bigcup_{P_{j}\in\mathbf{P}} P_{j} \right\}$$

to be the possible labels (PredOGs) assigned to the genes in $R_{i}$, which includes a label $P_{m}$ assigned to all missing genes for $R_{i}$. With this definition, we can obtain the probability of a gene in $R_{i}$ being classified into $P_{j}$

$$\mathbb{P}\left( P_{j} | R_{i} \right)=\frac{\left| R_{i}\bigcap P_{j} \right|}{\left| R_{i} \right|}$$

and $P_{m}$,

$$\mathbb{P}\left( P_{m} | R_{i} \right)=\frac{\left| P_{m} \right|}{\left| R_{i} \right|}$$

where

$$\left| P_{m} \right|= \left| R_{i} \right|-\sum_{P_{j}\in\mathbf{P}} \left| R_{i}\bigcap P_{j} \right|$$

According to the Shannon entropy, the entropy of a RefOG $R_{i}$ can be defined as

$$H_{i}\mathbb{=-P}\left( P_{m} | R_{i} \right)\log_{2} \mathbb{P}\left( P_{m} | R_{i} \right)-\sum_{P_{j}\in\mathbf{P}\left| R_{i}\bigcap P_{j}\neq\emptyset\right.} \mathbb{P}\left( P_{j} | R_{i} \right)\log_{2} \mathbb{P}\left( P_{j} | R_{i} \right)$$

Here, $H_{i}$ is unbounded, we can normalize it by dividing it by the maximum possible entropy a RefOG $R_{i}$ can achieve. The maximum entropy occurs when all genes in $R_{i}$ are equally distributed across all possible classifications.

$$NH_{i}=\frac{H_{i}}{\log_{2} \left| R_{i} \right|}$$

Here, the normalised entropy for each RefOG $R_{i}$ is denoted by $NH_{i}$, which is bounded between 0 and 1. If and only if all genes in $R_{i}$ are assigned to a single classification, $NH_{i}=0$.

Similar to recall and precision, we can compute the weighted average of normalised entropy across all RefOGs, weighted by the size of each RefOG $\left| R_{i} \right|$. This is given by,

$$\mathbf{H}=\frac{\sum_{i=0}^{S} \left| R_{i} \right|*{NH}_{i}}{\sum_{i=0}^{S} \left| R_{i} \right|}$$

$\mathbf{H}$ has the same interpretation as $NH_{i}$, but aggregated over all RefOGs. A small value indicates better performance, as it reflects lower uncertainty or more concentrated classifications for RefOGs, whereas a higher value indicates worse performance, as it reflects higher uncertainty or more scattered classifications.

The interpretation is similar to $NH_{i}$. The smaller the value of $\mathbf{H}$ is, the better the methods are.

8. Rank Score

Summary: Rank Score quantifies the overall performance of a method by ranking it relative to others on each evaluation measure, then averaging those ranks across all measures — lower scores indicate better overall performance.

To quantify the performance of each method, we introduce the rank score in this study, which takes the rank of each method against each measure mentioned above, then average the rank for each method across all measures.

Let $X$ be a random variable that takes elements from a countable set of methods $\mathcal{X=}\left\{ x_{1}, x_{2}, \ldots x_{q} \right\}, q\mathbb{\in N}$ if the number of methods. We can define a score function $s:X\mathbb{\to R}$ , such that $\forall x_{i}\mathcal{\in X,}s\left( x_{i} \right)\mathbb{\in R}$.

With the score function, a rank function $\rho:X\to\left\{ 1, 2, \ldots q \right\}$ which assigns a rank to each element $x_{i}\mathcal{\in X}$ based on its position in the sorted order can be expressed as follows:

$$\rho\left( x_{i} \right)=\left| \left\{ x_{j}\mathcal{\in X} | s\left( x_{j} \right)\leq s\left( x_{i} \right) \right\} \right|+1$$

here $\rho\left( x_{i} \right)$ counts the number of elements $x_{j}$ with scores less than $s\left( x_{i} \right)$ then adds 1 to give the rank.

Now let $\mathcal{M=}\left\{ m_{1}, m_{2}, \ldots, m_{p} \right\}$ be a countable set of measures, we can define the rank score for each method $x_{i}$ against each measure $m_{i}$ as

$$rs\left( x_{i} \right)=\left\{ \rho\left( x_{i} | m_{j} \right) | m_{j}\mathcal{\in M} \right\}$$

where $\rho\left( x_{i} | m_{i} \right)$ represents the rank of $x_{i}$ based on the measure $m_{j}$.

The final average rank score for method $x_{i}$ can then be given by

$$RS\left( x_{i} \right)=\frac{1}{p}\sum_{j=0}^{p} \rho\left( x_{i} | m_{j} \right)$$

where $p=\left| \mathcal{M} \right|$ is the number of measures.

In this study, we have seven measures and 13 methods, plug in those number of into the measure definition and the rank function we can obtain the table and figure shown below. They both illustrates the performance of each method using different measures. The last column in the table gives the average rank score for each method across all measures, so as the last plot in the figure.

Table 1. Methods scores for each Orthogroup benchmarking method across multiple tools.

| Methods | Missing RefOGs (%) | Missing Genes (%) | RefOG Fusions (%) | RefOG Fissions (%) | Recall | Precision | Entropy | F1-score | Rank Score |
| --- | --- | --- | --- | --- | --- | --- | --- | --- | --- |
| OF3_Align_ST | 0 | 2.533 | 2.857 | 32.857 | 84.112 | 82.94 | 0.097 | 78.492 | 2.625 |
| OF3_DB | 0 | 2.639 | 2.857 | 32.857 | 83.557 | 84.667 | 0.1 | 79.83 | 2.75 |
| OF3_Linear | 0 | 1.266 | 2.857 | 35.714 | 80.499 | 81.374 | 0.118 | 75.071 | 3.5 |
| OF2_DB | 0 | 2.533 | 2.857 | 35.714 | 78.686 | 83.268 | 0.131 | 74.524 | 3.625 |
| OF2_Align_ST | 0 | 2.533 | 2.857 | 35.714 | 78.686 | 83.268 | 0.131 | 74.524 | 3.625 |
| Broccoli | 0 | 3.113 | 2.857 | 27.143 | 75.572 | 80.425 | 0.131 | 70.912 | 4.875 |
| SP_sens | 0 | 5.805 | 2.857 | 35.714 | 70.81 | 85.864 | 0.18 | 68.911 | 5.625 |
| OrthoHMM | 0 | 3.166 | 2.857 | 31.429 | 76.183 | 75.725 | 0.137 | 66.395 | 6 |
| SP_def | 0 | 7.282 | 2.857 | 38.571 | 70.543 | 86.016 | 0.175 | 68.627 | 6.5 |
| OrthoMCL | 0 | 7.018 | 2.857 | 35.714 | 61.562 | 89.318 | 0.244 | 62.847 | 6.5 |
| SP_fast | 0 | 7.652 | 2.857 | 40 | 69.92 | 86.394 | 0.18 | 68.512 | 7 |
| ProteinOrtho | 0 | 16.201 | 2.857 | 65.714 | 37.355 | 95.814 | 0.363 | 47.508 | 9.125 |
| Hieranoid | 1.429 | 15.567 | 2.857 | 41.429 | 54.845 | 94.586 | 0.277 | 61.102 | 9.875 |
| FastOMA | 1.429 | 9.868 | 10 | 47.143 | 58.927 | 80.46 | 0.262 | 54.91 | 12.625 |
